# *DeepNeuron*: An Open Deep Learning Toolbox for Neuron Tracing

**DOI:** 10.1101/254318

**Authors:** Zhi Zhou, Hsien-Chi Kuo, Hanchuan Peng, Fuhui Long

## Abstract

Reconstructing three-dimensional (3D) morphology of neurons is essential to understanding brain structures and functions. Over the past decades, a number of neuron tracing tools including manual, semi-automatic, and fully automatic approaches have been developed to extract and analyze 3D neuronal structures. Nevertheless, most of them were developed based on coding certain rules to extract and connect structural components of a neuron, showing limited performance on complicated neuron morphology. Recently, deep learning outperforms many other machine learning methods in a wide range of image analysis and computer vision tasks. Here we developed a new open source toolbox, *DeepNeuron*, which uses deep learning networks to learn features and rules from data and trace neuron morphology in light microscopy images. *DeepNeuron* provides a family of modules to solve basic yet challenging problems in neuron tracing. These problems include but not limited to: (1) detecting neuron signal under different image conditions, (2) connecting neuronal signals into tree(s), (3) pruning and refining tree morphology, (4) quantifying the quality of morphology, and (5) classifying dendrites and axons in real time. We have tested *DeepNeuron* using light microscopy images including bright-field and confocal images of human and mouse brain, on which *DeepNeuron* demonstrates robustness and accuracy in neuron tracing.

## 1. Introduction

Over the past few decades, researchers have developed algorithms and tools to reconstruct (trace) 3D neuron morphology. A number of manual/semi-automatic neuron tracing software packages in both the public domain and commercial world have been developed (Meijering, et al., 2004; Peng, et al., 2010; Donohue and Ascoli, 2011; Longair, et al., 2011; Luisi, et al., 2011; Choromanska, et al., 2012; Peng, et al., 2014; Feng, et al., 2015). To further promote the development of neuron tracing tools, the DIADEM challenge (Liu, 2011) and the BigNeuron project (Peng, et al., 2015) were launched to compare different automated algorithms. At small or medium scales, many algorithms (base-tracers) have been shown to produce meaningful reconstructions on high quality neuron images. For large-scale image datasets, UltraTracer (Peng, et al., 2017) provides an extendible framework to scale up the capability of these base-tracers. Despite these efforts on algorithm and tool development, it remains an open question on how to faithfully reconstruct neuron morphology from challenging image datasets that have medium to low qualities and contain very complex neuron morphology.

Starting from a cell body, a neuron tracing process usually follows dendrites and axons, eventually connecting all such neuron signal as a tree that represents the morphology of the neuron. In light microscopy images, dendrites typically show continuous signal, whereas axons are often hard to trace due to their punctuated appearance and large, complex arborization patterns (Peng, et al, 2010s; see for example the bright-field images of biocytin-labeled neurons in the Allen Cell Type Database). In addition, the image quality varies a lot depending on sample preparation, imaging process, cell types and the healthiness of neurons. For instance, neuron signal could be continuous in one image, but dim and broken in another. It is difficult to automatically extract all such neuron signal under different conditions.

Several important steps in neuron tracing can be formulated as a classification problem. For example, detection of neuron signal from background is essentially foreground-background classification. Reconstruction of the topology of a neuron via connecting neuron fragments can be treated as connection-separation classification. In this aspect, a few studies used traditional machine learning and recent deep learning (LeCun, et al., 2015) models to produce neuron morphology. For example, Gala et al. introduced an active learning model by combining different features to automatically trace neurites (Gala, et al., 2014). Chen et al. proposed a self-learning based tracing approach, which did not require substantial human annotations (Chen, et al., 2015). Fakhry et al. (Fakhry, et al., 2016) and Li et al. (Li, et al., 2017) used deep learning neural networks to segment electron and light microscopy neuron images. Despite these algorithmic efforts, none of these methods provide publicly available tools to use on external datasets.

Nowadays deep learning methods outperform traditional methods in many pattern recognition and computer vision applications. We analyzed commonly used modules of neuron tracing/editing workflows in real applications, and concluded that an open source deep learning toolbox would help significantly to this growing field. Using deep learning neural networks as the classification models, we develop *DeepNeuron*, which provides several essential modules to neuron tracing. For automated tracing, *DeepNeuron* can be used as either a new tracing algorithm to reconstruct neurites from difficult neuron images, or an extra processing component to improve other tracing algorithms. *DeepNeuron* could also assist annotators in manual tracing. Supporting extendable functions as plugins, currently *DeepNeuron* contains five commonly used modules (Figure 1):

- **Neurite signal detection:** automatically identify 3D dendritic and axonal signal from background.
- **Neurite connection:** automatically connect local neurite signal to form neuronal trees.
- **Smart pruning:** filter false positive and refine automated reconstruction results.
- **Manual reconstruction evaluation:** evaluate manual reconstructions and provide quality scores.
- **Classification of dendrites and axons:** automatically classify neurite types during real-time annotation.

**Figure 1.**
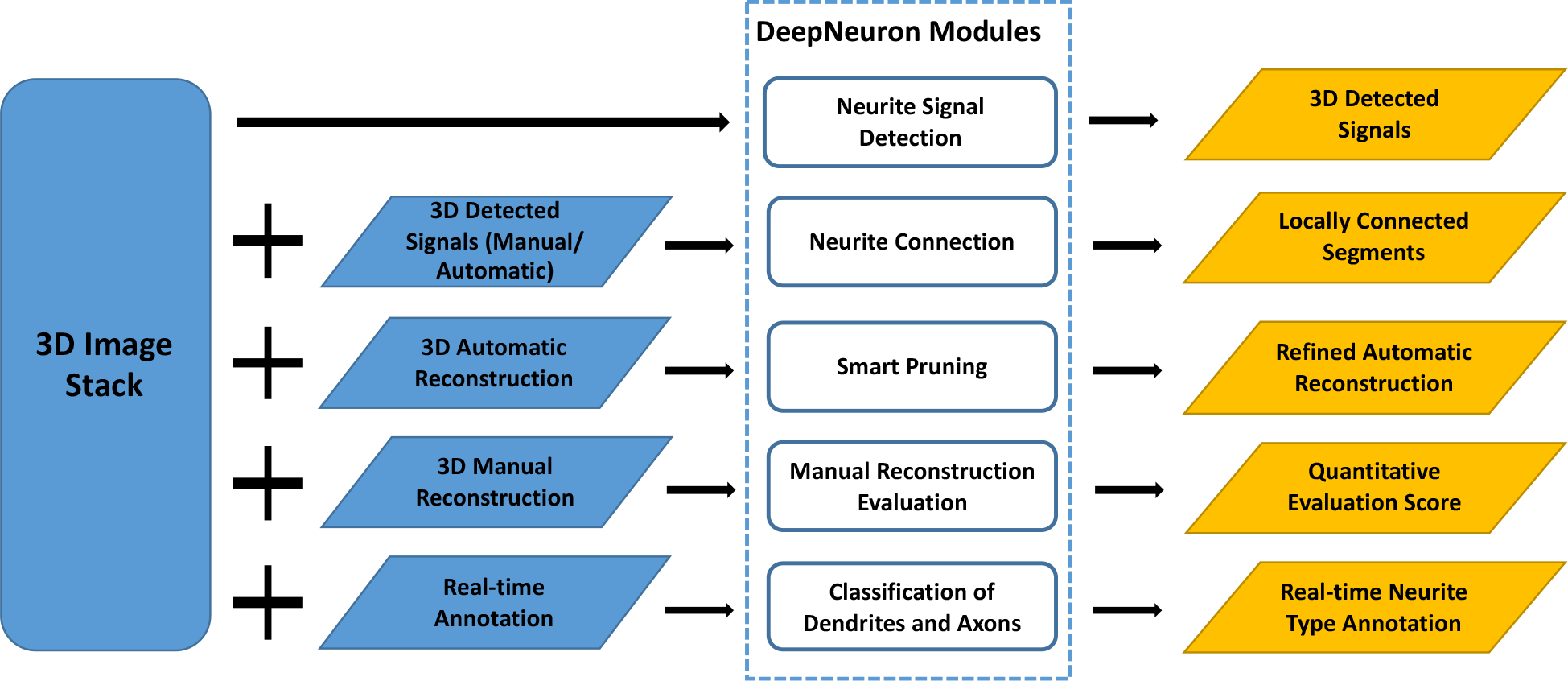
The workflow of the open source *DeepNeuron* toolbox, which has five deep learning based modules. Each *DeepNeuron* module has one or more processing components. Neurite signal detection module (section 2.1) uses convolutional neural networks (CNNs) to do foreground/background classification. Neurite connection module (section 2.2) uses a revised Siamese network (Bromley, et al., 1994; Chopra, et al., 2005) to connect neurite structure from detected neuron signals. Smart pruning module (section 2.3) refines a neuron’s morphology by using CNN models to filter out false positives. Manual reconstruction evaluation module (section 2.4) uses the output of CNNs as quality scores to evaluate reconstructions. Finally, dendrites/axons classification module (section 2.5) uses CNNs to perform multi-class classification to differentiate axons, dendrites, and background. Note that all the actual deep learning networks in our five modules can be replaced with other network models or user’s own design.

**Figure 2.**
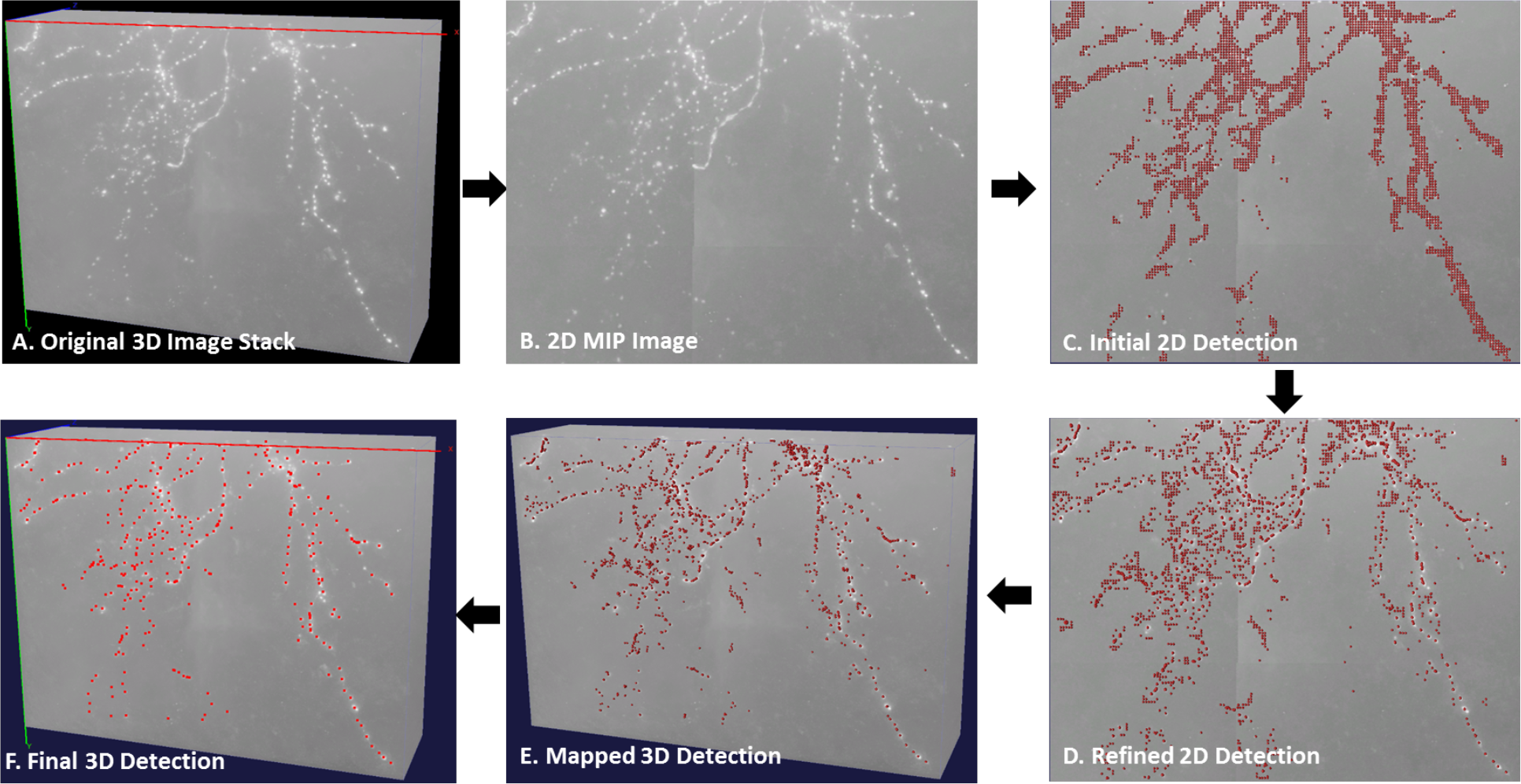
The workflow of 3D neurite signal detection. A) An example of the original 3D image stack. It is a cropped 3D bright-field image of a biocytin-labeled mouse neuron; the pixel resolution is 0.14 um x 0.14 um x 0.28 um. B) 2D MIP on the XY plane. C) Initial neurite signal detected by a deep CNN model (AlexNet in this case). D) Refined 2D signal detection result using a mean-shift. C and D are overlaid on top of B. E) Mapped 3D detection result based on local maximum intensity along Z-direction. F) Final 3D detection result after deep learning based refinement. E and F are overlaid on top of A. Red dots indicate 2D/3D detected signals.

## 2. Five Modules

### 2.1 Neurite Signal Detection

Due to difficulties in sample preparations and imaging, neurite signals often appear broken in a 3D image. It is hard to use any existing automated tracing algorithm to reconstruct 3D neuronal structures when this happens. Even for human annotators, locating these isolated axonal signals from the noisy background is a daunting work. To reliably detect neurite signals, we introduce the neurite signal detection module based on deep CNN to classify signal and background. This allows us to precisely detect neurite signals without any pre-processing steps applied on the original image. To speed up the detection and lower the GPU memory requirement, we used a two-dimensional (2D) CNN model followed by 3D mapping to detect signal in 3D and achieved satisfactory results on our testing data. However, our framework is not limited to 2D CNN but can also directly use 3D CNN models.

Manually reconstructed neurons were used as training samples. The 3D reconstruction of a neuron is represented as a tree, which contains a series of 3D X, Y, Z locations, radius, and topological “parent” of annotation nodes. To train the network, local 3D blocks (block size 61x 61 x 61 was used in our experiments) centered on manually annotated nodes in neurite segments were cropped from the original images. 2D maximum intensity projections (MIPs) of theses 3D blocks were used as the positive training set, and the same number of 2D background MIPs were randomly selected as the negative training set.

We tested our module using AlexNet (Krizhevsky, et al., 2012) with five convolutional and three fully-connected layers. Table 1 shows the five-fold cross-validation test of the module robustness. The training image dataset was partitioned into five equal size subsets (1-24, 25-48, 49-72, 73-96, and 97-122 as shown in Table 1). Four subsets were used for training, and the remaining single subset was used for validation. Our results show our overall accuracy > 98% for both training and validation.

**Table 1.** Five-fold cross-validation on bright-field training sets

| Training Set | Foreground Accuracy |  | Background Accuracy |  | Overall Accuracy |  |
| --- | --- | --- | --- | --- | --- | --- |
|  | Training | Validation | Training | Validation | Training | Validation |
| $\{1-122\} \setminus \{1-24\}$ | 98.77% | 97.78% | 98.97% | 96.87% | 98.87% | 97.33% |
| $\{1-122\} \setminus \{25-48\}$ | 98.54% | 99.07% | 98.78% | 98.34% | 98.66% | 98.71% |
| $\{1-122\} \setminus \{49-72\}$ | 98.67% | 98.28% | 98.78% | 99.13% | 98.73% | 98.71% |
| $\{1-122\} \setminus \{73-96\}$ | 98.71% | 96.64% | 98.28% | 99.23% | 98.50% | 97.94% |
| $\{1-122\} \setminus \{97-122\}$ | 98.64% | 99.02% | 98.77% | 98.41% | 98.71% | 98.72% |
| Average | **98.67%** | **98.08%** | **98.83%** | **98.44%** | **98.75%** | **98.26%** |

In testing, we first projected the original 3D image stack onto the XY plane and generated a MIP image. We then cropped 2D patches using a sliding window with *n*-pixel stride. These patches were classified into patches centered on foreground or background pixels using our trained CNN model. To further improve classification accuracy and exclude false positive patches, we applied mean-shift (Cheng, 1995) to the detected foreground patches and map them back to the actual 3D locations based on the local maximum intensity along Z. Finally, we classified these 3D detected signals using our CNN model again based on the MIPs of the local 3D blocks.

We applied our module to two challenging datasets of mouse neurons. The first set was a bright-field biocytin-labeled mouse neuron dataset from Allen Cell Type Database. The second set was a whole mouse brain data imaged by fMOST imaging technology (Gong, et al., 2016). For the first dataset, we used 122 bright-field neuron image stacks and their associated manual reconstructions as the training set and produced ~813K training samples including ~404K foreground patches, and ~408K background patches. For the whole mouse brain dataset, we used ~493K training samples including ~252K foreground patches, and ~241K background patches from 22 whole mouse brain images. Figure 3 shows two examples of axon detection results. Using neurite signal detection module, most of axonal signals have been precisely detected in both datasets.

**Figure 3.**
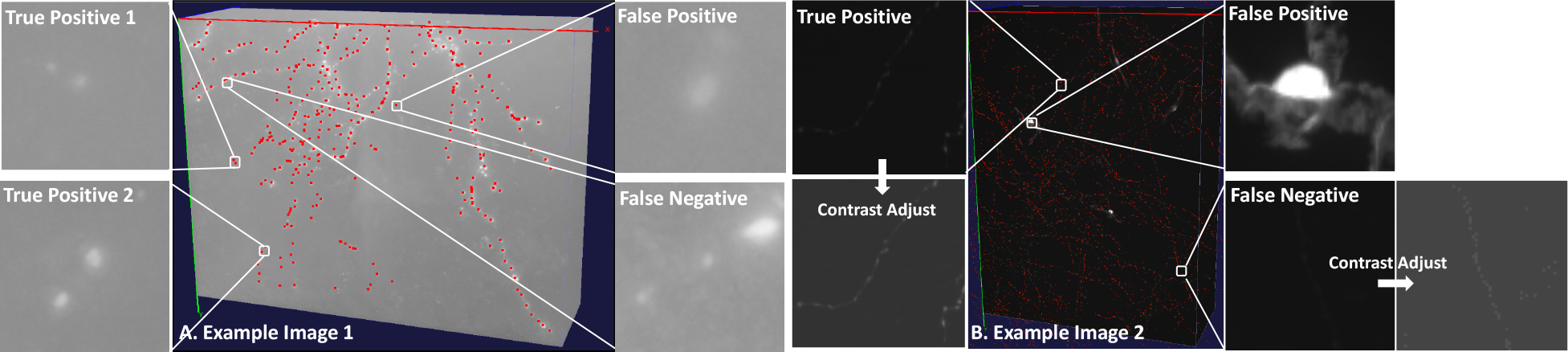
Axon detection results on two challenging datasets. A) An example of a 3D image stack of mouse neuron imaged with bright field microscopy (also see Figure 2A). B) An example of a 3D stack (shown as cropped) from a whole mouse brain imaged with fMOST; the pixel resolution is 0.3 um x 0.3 um x 1 um. Red dots indicate detected axonal signals. The two false positive example patches shown in A and B could be eliminated with more training samples. Those false negative patches (also as shown in A and B) with very weak signals at the center could be further identified by increasing the amount of weak signal foreground samples in the training set.

### 2.2 Neurite Connection

A complete neuron forms a tree structure that is composed of continuous neurite segments. Global, local, and topological features including total length, bifurcations, terminal tips and more others are used to study the neuronal morphology. These features have to be extracted from neurite segments instead of dots. Therefore, finding the continuity of neurites and connecting neurite segments is a critical step in neuron tracing. Generally, automated tracing algorithms can achieve good performance on connecting neurite segments with small gaps based on the continuity of segment orientations. However, it is difficult to automatically connect dots-like neurite signals. Using the spatial distance between these signals as the weight, Minimal Spanning Tree (MST) provides a possible solution. However, without biological context, it could also introduce topological errors. Human beings are good at finding the continuity of isolated signals per their observations and domain knowledge. By learning the neurite connectivity from a large dataset annotated by humans, a deep learning based MST (DMST) approach we proposed can successfully connect neurite segments with relatively big gaps (Figure 4).

**Figure 4.**
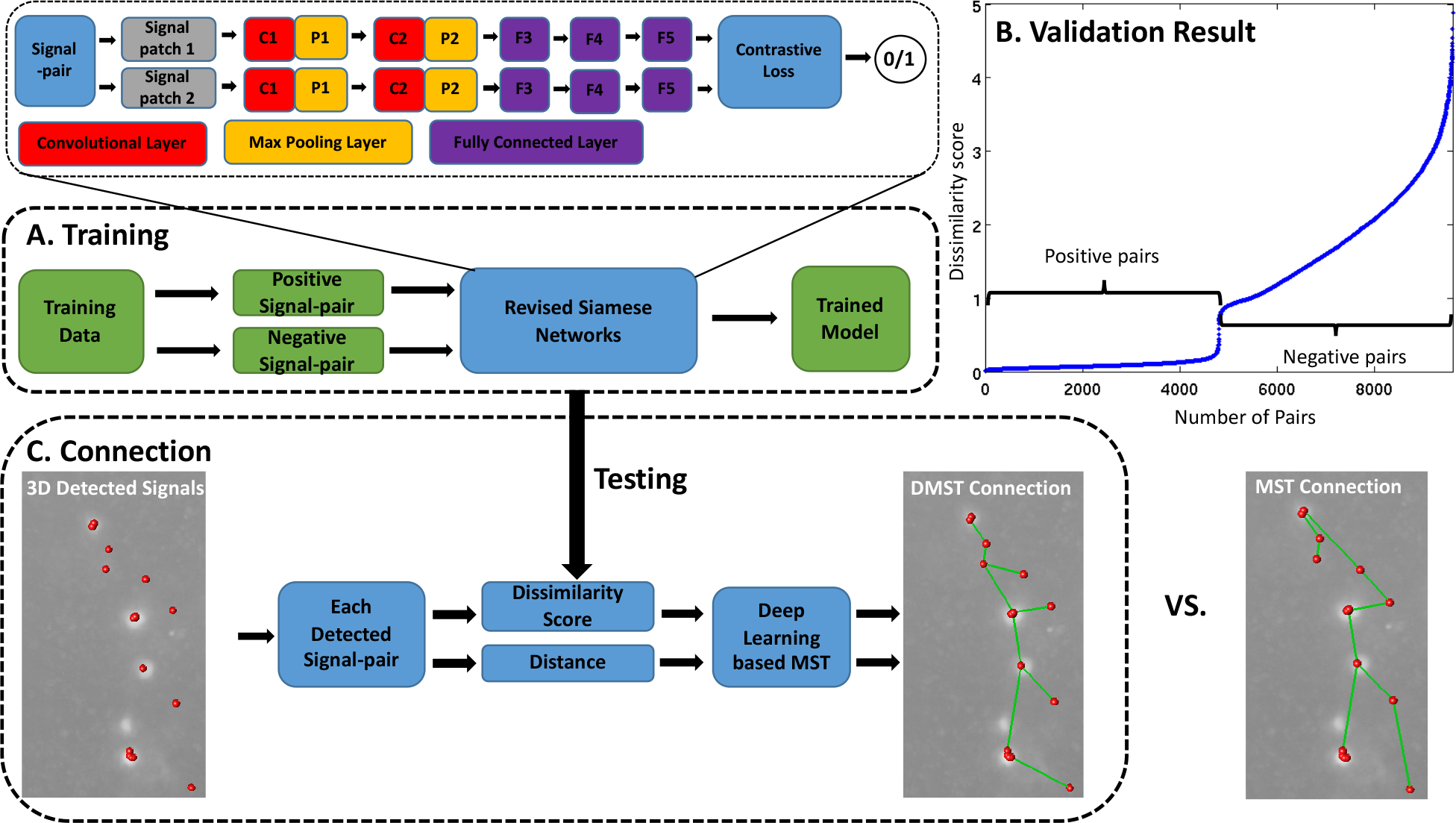
The workflow of neurite connection module. A) In the training step, the connectivity of a signal pair is learned using a revised Siamese network. In each pair, two 1×200 feature vector are extracted. B) The validated result with sorted dissimilarity scores for all signal pairs shows that the dissimilarity scores of positive pairs are much lower than that of negative pairs. C) The trained model is applied in the connection step to calculate the dissimilarity for each detected signal pair. Results of our DMST connection and the original MST connection (using distance as the weight only) are shown.

Siamese networks (Bromley, Guyon, LeCun, Säckinger and Shah, 1994; Chopra, Hadsell and LeCun, 2005) are used among tasks that involve finding similarity or the relationship between two subjects being compared. Our revised Siamese model in this work includes two identical arms. Each consists two convolutional layers with max pooling, followed by three fully-connected layers. The two arms are then fed to a contrastive loss function to produce a binary decision.

In training (Figure 4A), we used pairs of patches generated from two consecutive annotation nodes as positive training samples, and pairs of patches generated from two spatially separated annotation nodes as negative training samples. We used ~919K training pairs, ~460K of them being positive pairs and ~459K being negative pairs.

In connection (Figure 4C), a 1xM feature vector is extracted from individual input patch (M can be defined by the user. We used M = 200 in our experiment). The Euclidean distance between two feature vectors is calculated as the dissimilarity score of a patch-pair, which is multiplied by the distance to form the weight in our proposed DMST graph.

Combing neurite signal detection and neurite connection modules, we were able to reconstruct axons that present big challenges to traditional methods due to large gaps between signal segments (Figure 5).

**Figure 5.**
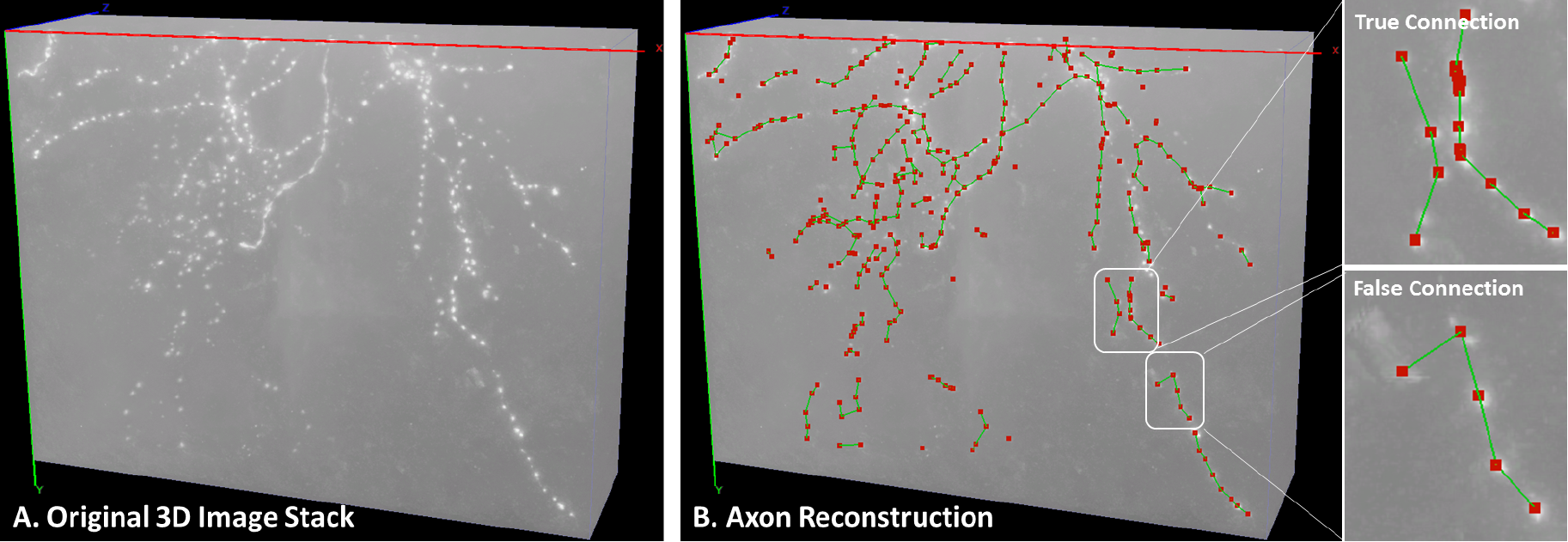
*DeepNeuron* axon reconstruction. A) The same example image as shown in Figure 2A. B) 3D axon signals (red dots) were extracted with neurite signal detection module; local 3D connections (green lines) between signals were generated by neurite connection module. As shown in B, the proposed DMST in the connection module can formulate the correct neurite structure even when the detected signals from different neurites are spatially close to one another. In a more difficult case, however, the loss of true signal in the local area can make the information captured by the network outweighed by the distance, resulting in false connection. This could be avoided by adequately enlarging the examined local area. Note that unconnected fragments like those on the bottom-left would be further processed by our non-deep-learning functions of the tool, which is out of the scope of this article.

### 2.3 Smart Pruning

Many of the existing automatic tracing algorithms rely on the correct estimation of the threshold that separates the potential foreground signal from the background. Typical methods include those that use the weighted average intensity of the entire image to threshold the image or add a pre-processing step to enhance signals. These methods have limited success for neuron images with low signal-to-noise ratio and uneven background. To solve the problem, we developed a smart pruning module in *DeepNeuron*. It relieves the burden of precisely separating foreground and background up the front. Using existing algorithms, our module first generated over-traced results with a lower foreground/background segregation threshold or through signal enhancement step. We then trained CNN networks to classify true signals and false positive signals. Using the trained models, we filtered out falsely detected signals and pruned the reconstructed neuronal tree. Furthermore, different tracing results generated from multiple base tracing algorithms could be combined (Wan, et al., 2015) to produce a consensus using this module (Figure 6).

**Figure 6.**
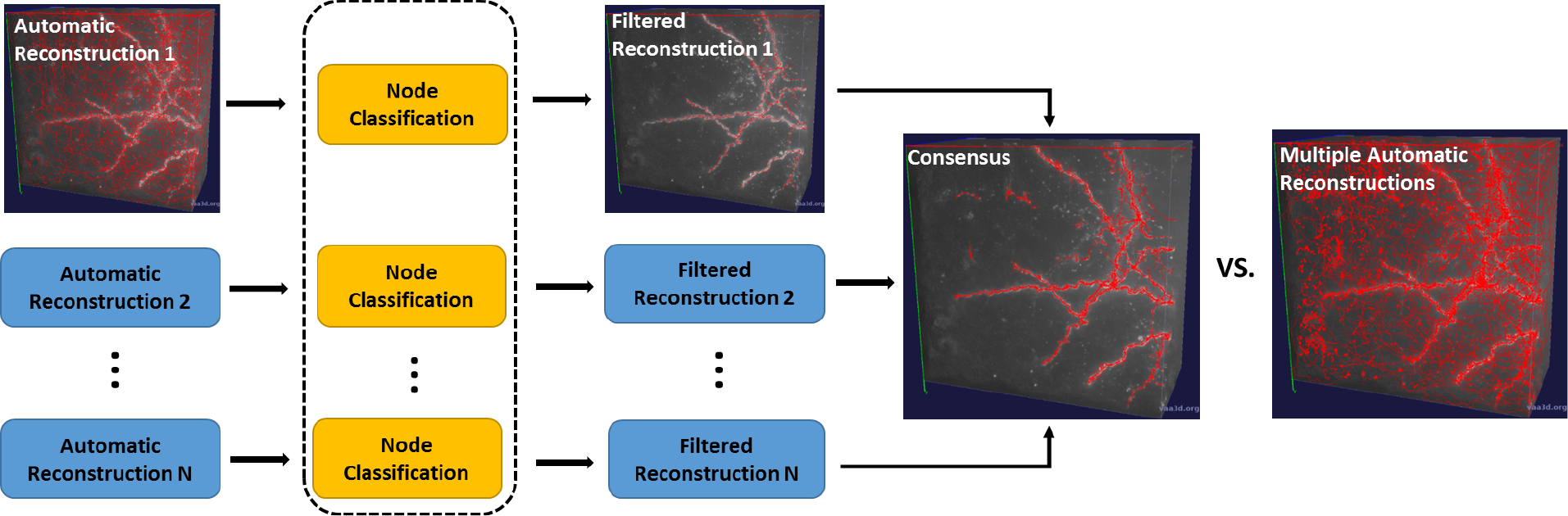
The workflow of consensus generation using the smart pruning module. Multiple automatic reconstructions are filtered by CNN based classification models first. Then all filtered reconstructions are fused together to produce a consensus. Reconstructions are shown in red lines on top of the original image stack.

### 2.4 Manual Reconstruction Evaluation

Since manual reconstruction is largely used as the gold standard to evaluate automated reconstruction algorithms and to generate training set for machine learning based approaches, it is important to assess the consistency of manual reconstructions among different annotators or of the same annotator at different times. For this purpose, *DeepNeuron* provides an evaluation module based on deep learning classification model. Take the mouse neuron dataset from the Allen Cell Type Database we described in section 2.1 as an example, we divided the 122 manual reconstructions from multiple annotators into five subsets and took a five-fold cross validation strategy. Each time we took four subsets as the training data. Once the network was trained, we used it to evaluate how consistent the remaining subset is with respect to the training subsets (Figure 7, Table 2). More specifically:

- First, all annotation nodes in test subset were classified into two categories: foreground and background.
- All classified foreground nodes formed an initial prediction.
- Based on the orientation, tip location, and distance, fragments in the initial prediction were automatically connected to produce a refined prediction. In our experiment, we only connected terminal tips between two segments whose orientation differs less than 30 degrees and distance is smaller than 30 voxels.
- The test subset is evaluated by the consistency score *c*:

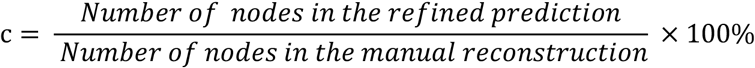

**Figure 7.**
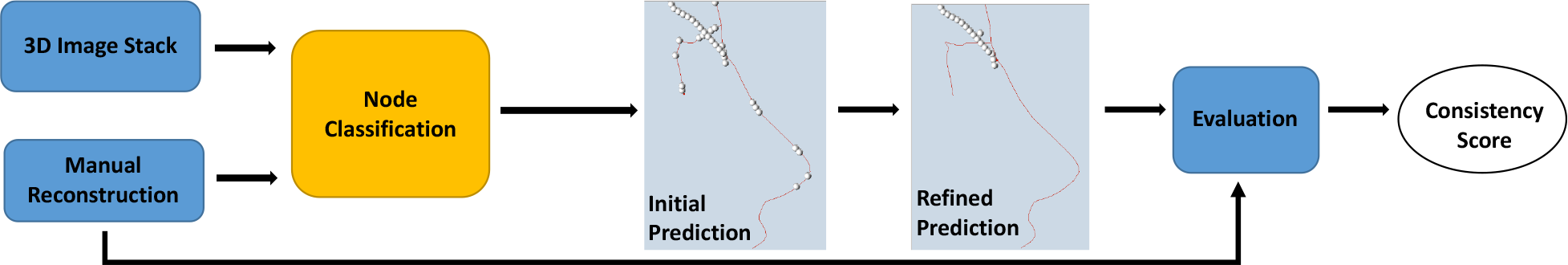
The workflow of manual reconstruction evaluation module. All annotation nodes are classified into foreground (red lines) and background (white dots). Gaps in the initial prediction were automatically filled based on the orientation, tip location, and distance in the refined prediction.

**Table 2.**
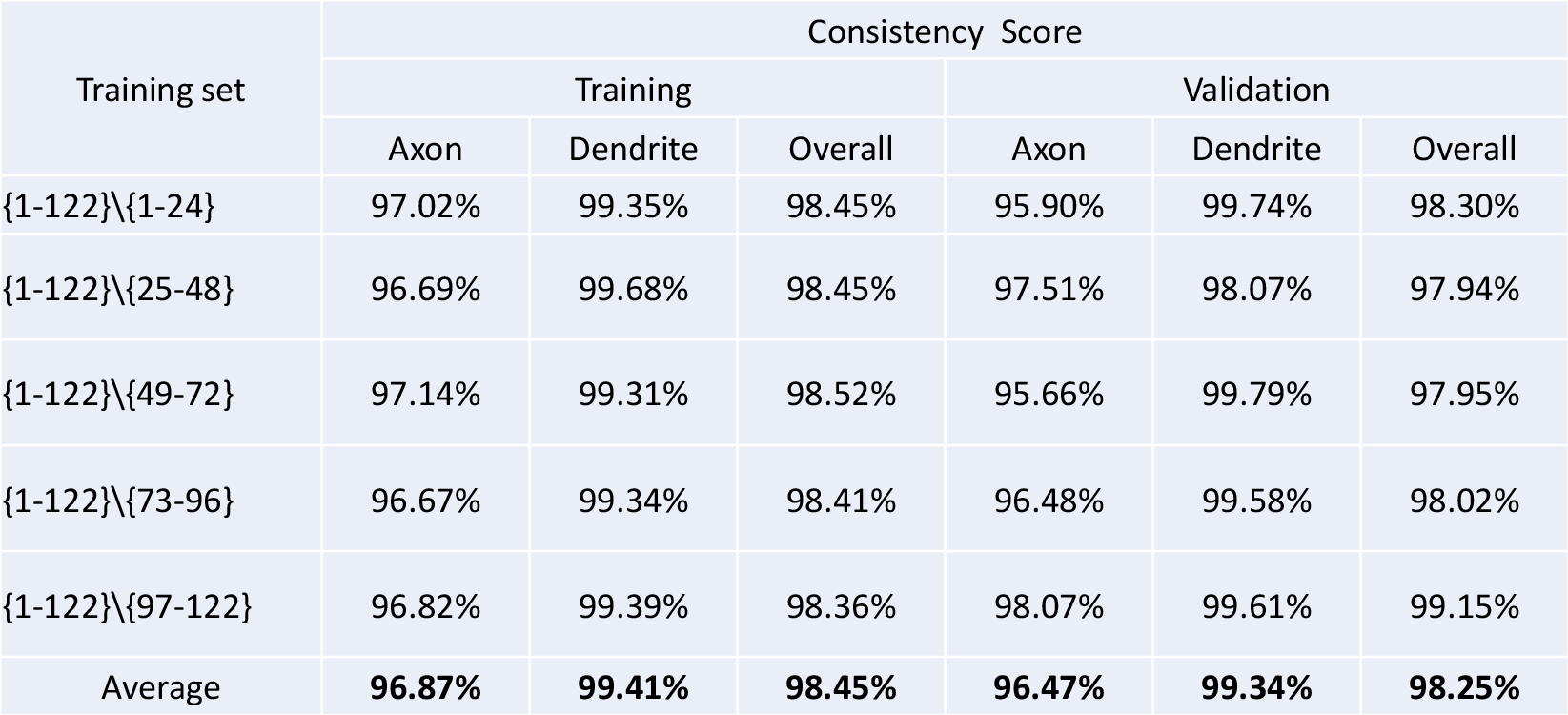
Five-fold cross-validation on 122 manual reconstructions of biocytin-labeled mouse neuron dataset

Table 2 shows the five-fold cross validation results on 122 manual reconstructions for the bright-field biocytin-labeled mouse neuron dataset from Allen Cell Type Database. The high consistency scores indicate that manual reconstructions are very consistent across different annotators and different subsets of data. In addition, thicker and more continuous dendrites (>99%) have higher consistency scores than dim and discontinuous axons (>96%), which are harder to reconstruct.

More broadly, we applied our evaluation module to 31 manual reconstructions including 10 human neurons and 21 mouse neurons in the Allen Cell Type Database. Table 3 shows our comparison results. Consistent with Table 2, dendrites have higher scores than axons. In addition, scores on human neurite (axon and dendrite) reconstructions are higher than those of mouse, indicating that annotators have better tracing performance on physically larger human neurons.

**Table 3.** Comparison results of consistency scores on human and mouse neuron reconstructions

|  | Number | Axon | Dendrite | Overall |
| --- | --- | --- | --- | --- |
| Human | 10 | 98.23% | 99.60% | 98.93% |
| Mouse | 21 | 94.28% | 99.40% | 95.38% |

### 2.5 Classification of Dendrites and Axons

Dendrites and axons have their own functions and play different roles in the nervous system. Distinguishing these two types of neurites can help us gain insight into the brain circuitry. Although dendrites and axons show different shapes and intensity properties in light microscopy images, such a general rule of thumb, however, is not always guaranteed. Due to variant image quality, axons can also appear continuous and look more like dendrites. This makes them difficult to be correctly labeled in most of tracing algorithms. Here we present a deep learning module serving as a vehicle for the networks that are trained for this purpose. This tool allows to automatically classify dendrites and axons on real-time manual annotation, and potentially save time for annotators.

We used the same approach as described in section 2.1, except that the problem is now a multinomial classification (dendrite, axon, and background) instead of a binary classification (foreground and background) problem. Figure 8 shows the performance on two testing cases from the Allen Cell Type Database. In this example, we used ~813K training samples including ~143K axons, ~261K dendrites, and ~409K background. In this article, AlexNet and a revised model were used as demonstration. In Fig. 8, both AlexNet and our revised model can accurately classify continuous dendritic and discrete axonal signals. However, when the signal of axons is similar to dendrites (Figure 8B), AlexNet mistakenly classified axons into dendrites, while the revised model with one more convolutional layer successfully distinguished axons from dendrites. Table 4 shows the comparison of classification accuracy between the two models on the training samples. We found that the revised model yields much better classification performance. In exchange, the efficiency of the revised model is sacrificed due to much more number of outputs in each convolutional layer (3.27s forward-backward time for AlexNet; 335.91s forward-backward time for the revised model).

**Figure 8.**
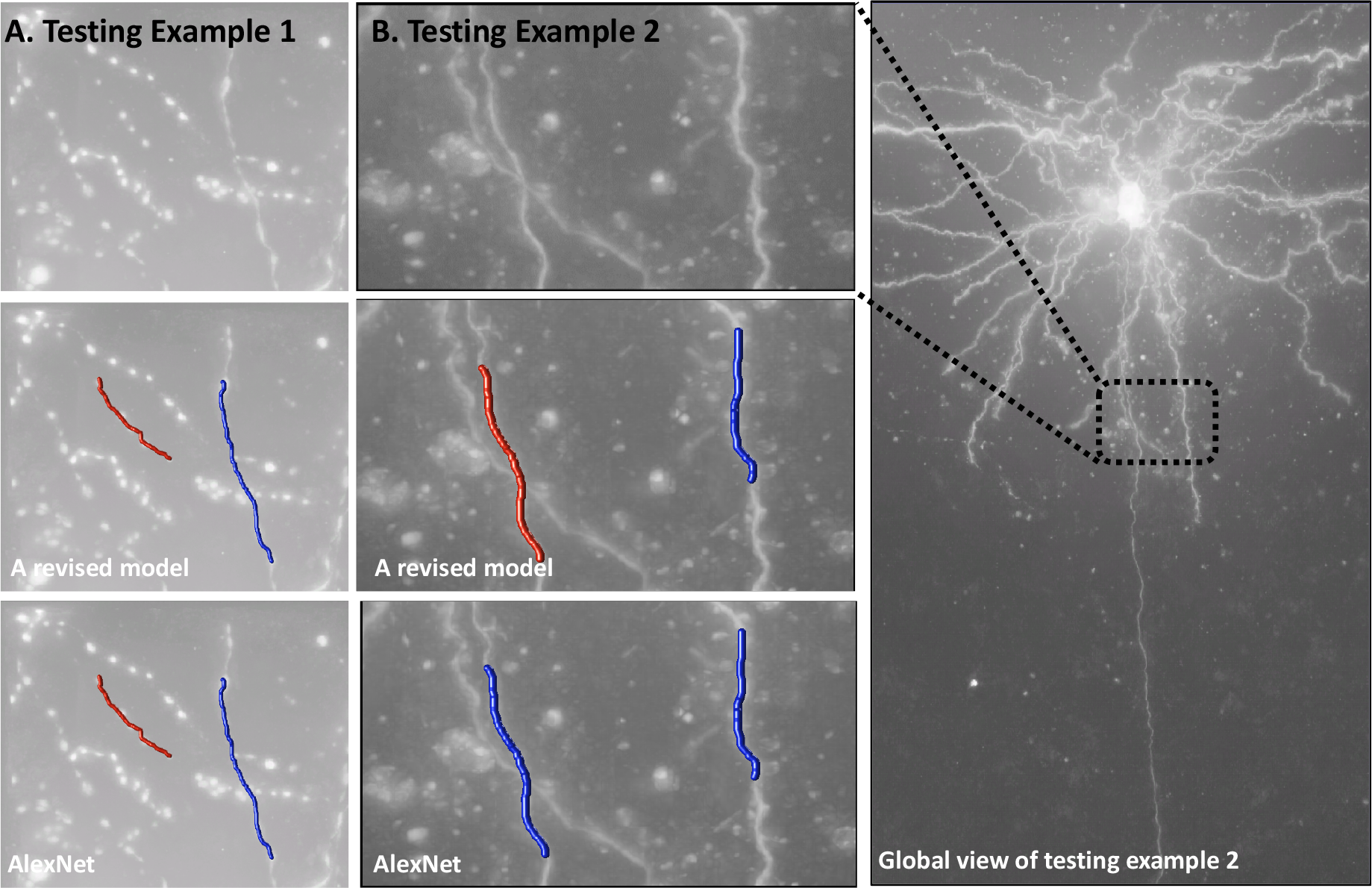
Comparison of dendrite and axon classification using AlexNet and a revised model. A) Axons are discrete, and dendrites are continuous. B) Both axons and dendrites are continuous. The left segment is extracted from a long axon. The right segment is extracted from a local dendrite. Red color indicates the axon, and blue color indicates the dendrite in A and B. Note that all these 3D segments are manually annotated using Virtual Finger technology (Peng, et al., 2014); neurite types are automatically annotated by the proposed module.

**Table 4.** Comparing AlexNet with a revised model

| Deep learning network | Averaged forward-backward time | Deemed axon | Deemed dendrite | Deemed background |
| --- | --- | --- | --- | --- |
| AlexNet | 3.27 s | 84.79% | 93.86% | 99.04% |
| A revised model | 335.91 s | 97.89% | 98.55% | 99.82% |

## 3. Discussion

In this paper, we presented a new deep learning based open source toolbox for neuron tracing: *DeepNeuron*. With extensible framework, *DeepNeuron* currently provides five modules to comprehend the major tasks:

- For a neuron image stack, it can be used to automatically detect neurite signals.
- For a neuron image stack with detected 3D signals, it can automatically connect signals to generate local segments.
- For a neuron image stack with its associated automated reconstruction, it can be used as a filter to clean up all false positive tracing and generate a refined result.
- For a neuron image stack with its associated manual reconstructions, it can evaluate how consistent and reliable the reconstructions are.
- For a neuron image stack with interactive human annotation via the user interface, it can label neurite types in real-time.

*DeepNeuron* has been implemented as an open source plugin in Vaa3D (http://vaa3d.org) (Peng, Ruan, Long, Simpson and Myers, 2010; Peng, Bria, Zhou, Iannello and Long, 2014). *DeepNeuron* toolbox is a highly flexible vehicle allowing investigators to take advantage of deep learning to facilitate neuron tracing in their research. As mentioned in this article, researchers can freely replace different network models that suit their needs. Combined with other related features in Vaa3D including 30+ automatic neuron tracing plugins, semi-automatic neuron annotation, annotation utilities, neuron image/reconstruction visualization, *DeepNeuron* works as a smart artificial intelligence engine which offers great help to biologists in exploring neuronal morphology.

## Toolbox and Software Availability

The *DeepNeuron* toolbox was written in C++ as a plugin to Vaa3D. *DeepNeuron* source code is available at https://github.com/Vaa3D/vaa3d_tools/tree/master/hackathon/MK/DeepNeuron. In addition, the *DeepNeuron* plugin is also included as a plugin in binary releases of Vaa3D, which can be downloaded at https://github.com/Vaa3D/Vaa3D_Data/releases/tag/1.0.

## Acknowledgement

We thank Allen Institute for Brain Science for providing neuron datasets and manual annotations. The authors wish to thank the Allen Institute founders, P. G. Allen and J. Allen, for their vision, encouragement and support.

## Author contributions

H.P. conceived and managed the project. F.L. proposed the overall technical framework. Z.Z. developed the toolbox and conducted the experiments. H.K. implemented a plugin for the dendrite and axon classification and assisted in several other experiments. All authors edited the manuscript.

## Conflict of Interest Statement

On behalf of all authors, the corresponding author states that there is no conflict of interest.

